# Dynamical effects of dendritic pruning implicated in aging and neurodegeneration: Towards a measure of neuronal reserve

**DOI:** 10.1101/2020.04.09.035048

**Authors:** Christoph Kirch, Leonardo L. Gollo

## Abstract

Aging is a main risk factor for neurodegenerative disorders including Alzheimer’s disease. It is often accompanied by reduced cognitive functions, gray-matter volume, and dendritic integrity. Although age-related brain structural changes have been observed across multiple scales, their functional implications remain largely unknown. Here we simulate the aging effects on neuronal morphology as dendritic pruning and characterize its dynamical implications. Utilizing a minimal computational modeling approach, we simulate the dynamics of detailed digitally reconstructed pyramidal neurons of humans obtained from the online repository Neuromorpho.org. We show that as aging progressively affects neuronal integrity, neuronal firing rate is reduced, which causes a reduction in energy consumption, energy efficiency, and dynamic range. Pruned neurons require less energy but their function is often impaired, which can explain the diminished ability to distinguish between similar experiences (pattern separation) in older people. Our measures indicate that the resilience of neuronal dynamics is neuron-specific, heterogeneous, and strongly affected by dendritic topology and the centrality of the soma. Based on the emergent neuronal dynamics, we propose to classify the effects of dendritic deterioration, and put forward that soma centrality measures neuronal reserve. Moreover, our findings suggest that increasing dendritic excitability could partially mitigate the dynamical effects of aging.

## Introduction

Aging is often accompanied by cognitive deficits such as forgetfulness, distractibility, inflexibility [1], reduction in speed of processing [2], working and episodic memory [3], inhibitory function [4], and long-term memory. Along with these cognitive changes, aging is also a major risk factor for dementia and neurodegenerative disorders [5]. There are a number of age-related structural changes that occur in the brain at various levels: synapses [6], dendrites [7], circuits [8], and gray-matter volume [9]. Furthermore, some dynamic properties associated with aging have been reported [10], however, the effect of dendritic pruning caused by aging on neuronal dynamics remains largely unknown.

It is well established that neurons can undergo several major structural changes during aging [11, 12]. Firstly, the number of dendritic spines decreases [6, 11]. As spines are regions of dense synaptic connections, neurons with fewer spines are often subjected to a reduced number of incoming synapses. Secondly, aging can alter the geometry of dendritic branches. For example, it has been found that aging neurons in humans show lumps and localized swelling that were not present in the control group [13]. Thirdly, the complexity and size of the dendritic arbor is progressively reduced. In pyramidal neurons, a recession of basal dendrites has been observed, followed by a retraction of the apical dendrites towards the main shaft [13]. Over time, the protrusion and branching order of neurons can decrease significantly, though the exact timescales over which these changes occur are not well known. These morphological changes have been observed across species [14-17] and in different cell types [15, 18, 19], with varying severity. The effect of aging is amplified by high senility [13], or in the presence of neurodegenerative disorders [11, 20, 21].

Given that dendrites perform vital computations that shape the neuron’s output [22-24], changing the structure can impede normal neuronal functioning and disrupt neural circuits. Exactly what adverse effects do a receding dendritic tree induce? What types of structural changes are detrimental? These questions cannot be answered easily because of the complex biophysical behavior of neurons that depends on a large number of dynamic internal and external factors. While associations of the effects of aging neurons exist – such as memory loss [1], a decrease in cognitive ability [25], and lower reaction and processing speeds [2] – the specific role that topology, and changes therein, plays in the functioning of single neurons is not well known [6, 26]. Furthermore, a number of factors (years of education, lifestyle, occupation, genetics, brain size, number of synapses, and so on) have been proposed to explain the broad diversity that is observed in the impact of cognitive impairments associated with neurodegeneration. Identifying the importance of some of these factors has been a main approach to explain differences in resilience to neurodegenerative damage within a population. Within this realm, the accumulation of protective factors increases the resilience. This is the foundation of influential concepts like brain reserve [27], and cognitive reserve [28]. Nonetheless, the neuronal basis of these protective factors is unclear.

Previous work proposed to apply a minimal model to the dynamics of active dendrites and to explore the behavior of realistic neuron morphologies that have been stored in an open database with over 100,000 neurons [29]. In this way, the dendritic structure of the neuronal modes can be extremely detailed and realistic, and provide insights into the spatial dependence of the dynamics. In this paper, we will extend the model to address: How do topological changes, as seen in aging and neurodegenerative disorders, impede a neuron’s normal functioning? To this end, we develop a simple pruning algorithm (available at http://www.sng.org.au/Downloads) that models the effects of aging on neuronal structure. Further, we characterize fundamental dynamical properties (energy consumption and dynamic range) of human pyramidal neurons receiving stochastic synaptic input at hundreds to thousands of compartments as the levels of neuronal degeneration (dendritic pruning) progressively accumulate.

## Results

To model the dendritic regression often observed in aging neurons, we iteratively removed the terminal compartments of the neuron reconstructions (see Methods for details). This pruning algorithm reproduces a reduction in the size and complexity of the dendritic arbor (Fig. 1). In this example of a human pyramidal neuron, the basal dendrites slowly disappear, while the apical tuft retracts towards the main stem. A critical change occurs in the topology at iteration 24, at which the number of dendritic stems connecting to the soma reduces to 1. The relative centrality of the soma reaches 0 in the previous iteration step because, topologically, it is interchangeable with the terminal compartment that makes up the branch that disappears in the next pruning iteration. After the critical point, only a single bifurcation remains at the top of the apical dendrites. It disappears at iteration 27. After this, the stem linearly reduces in size until disappearing completely at iteration 46.

**Figure 1:**
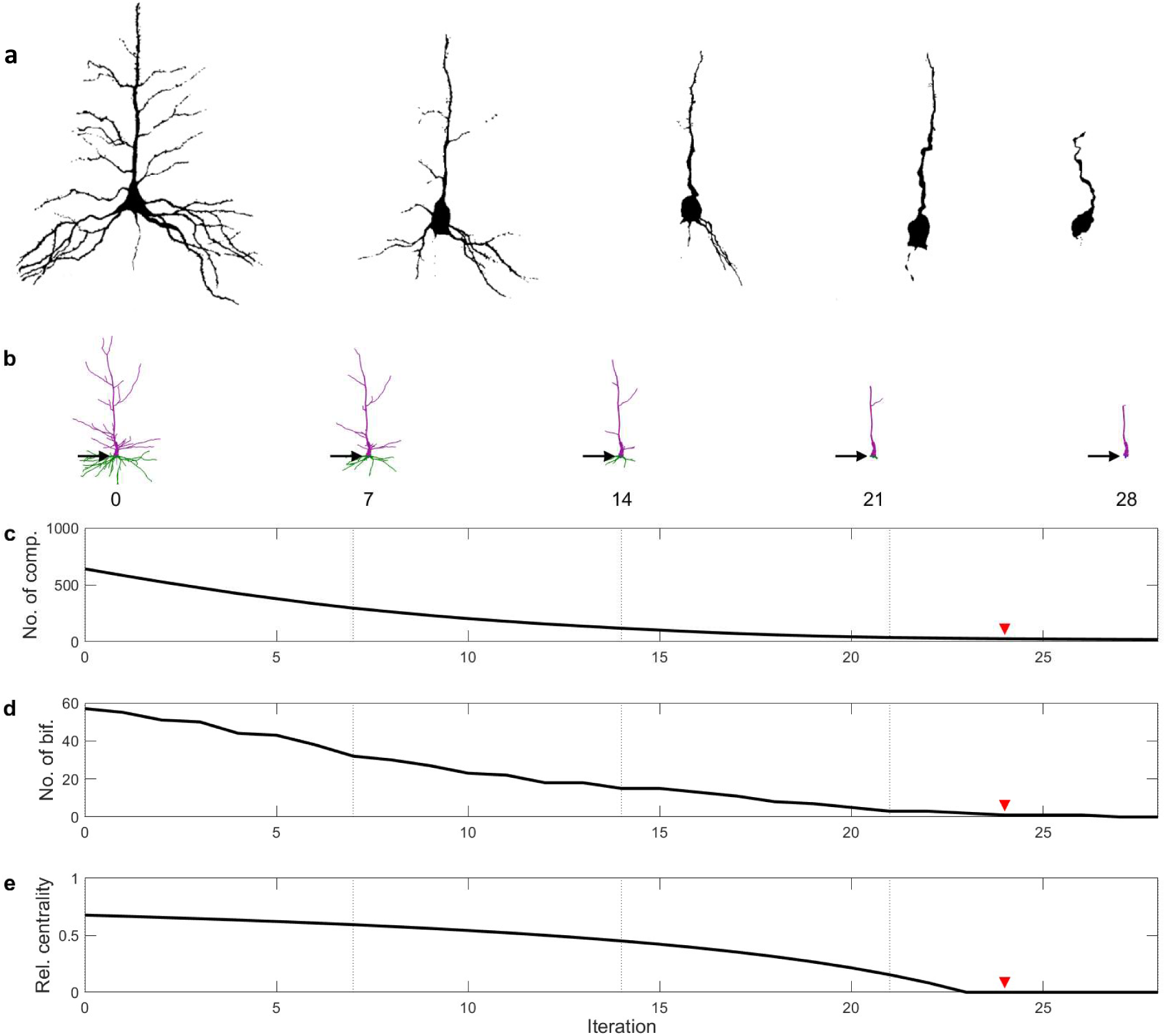
Dendritic pruning and neuronal aging. (a) Human neurons in their various aging stages (adapted with permission from [13]). (b) Snapshots of a digitally recreated human neuron (Neuron A, see Methods) in various stages of simulated aging. The number below each neuron signifies the pruning iteration (see Methods). Basal dendrites are colored in green, while apical dendrites are colored purple. The soma (blue) is shown by the black arrow. (c) The number of compartments as a function of the pruning iteration. (d) The number of bifurcations as a function of the pruning iteration. (e) Relative centrality of the pruned neuron (see Methods). At the red arrow, a single dendritic branch remains, which bifurcates once at the top of the apical tree.

For the different pruning iterations, we simulated the dynamics of the neuron, considering the dynamics of each compartment as a minimal excitable system (modeled as a cyclic cellular automaton and illustrated in Figs. 2 a, b). We studied six neurons (labelled Neuron A, B, …, F; see methods for details). As explained in detail in the Methods, each quiescent compartment can become active driven by stochastic synapses with a rate *h*, or by a propagation of input from a neighbor compartment with probability *P*. For a full illustration of the dynamics of the model, see Supplementary Video 1. A snapshot of the video is presented in Fig. 2 c. As demonstrated previously [29], the average firing rate is heterogeneous and depends on the neuronal topology (Fig. 2d), which may facilitate dendritic computations [23]. For each compartment, the firing rate and dynamic range can be measured, and they show specific spatial patterns such as amplification of dynamic range at bifurcation points [29]. Here we will focus on these measures specifically at the soma (Fig. 2 e). In addition, we will also focus on the total energy consumption of neurons and the relative energy consumption of the neuron (as a function of the two control parameters: *P* and *h*, and shown in Figs. 2 f, g). Here, the total energy consumption is defined as the average number of total dendritic spikes observed for every one somatic spike. The relative energy consumption indicates the mean number of dendritic spikes across dendritic compartments observed for each somatic spike (see Methods). As demonstrated in Fig. 2h, the relative energy consumption was previously shown to be an effective index to classify different types of neuronal dynamics [29]. This dynamical classification reflects two topological features of dendrites: centrality of the soma, and the number of dendritic branches connected to the soma. These dynamical features shown in Fig. 2 will be explored in more detail here as neuronal dendrites are progressively pruned.

**Figure 2:**
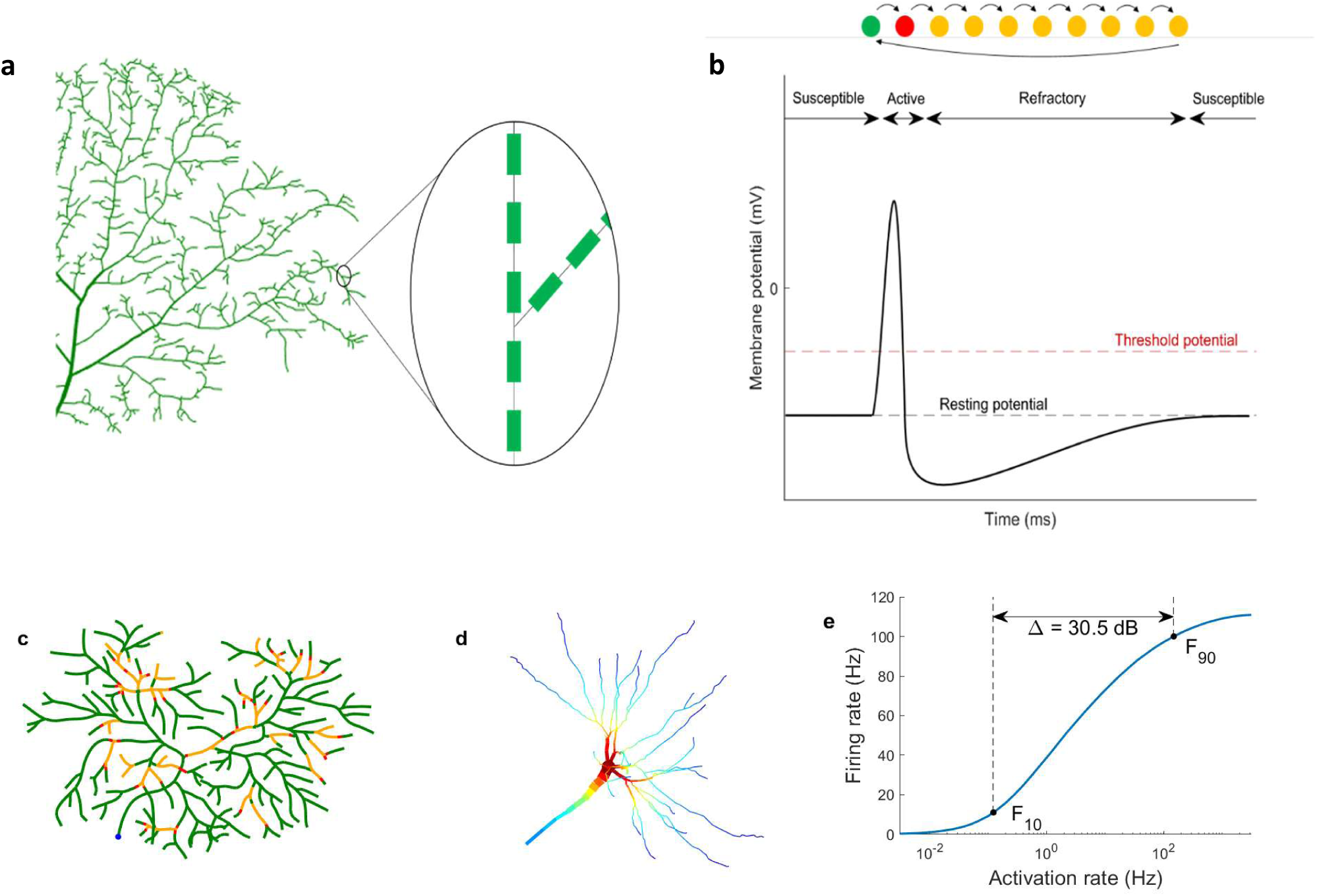
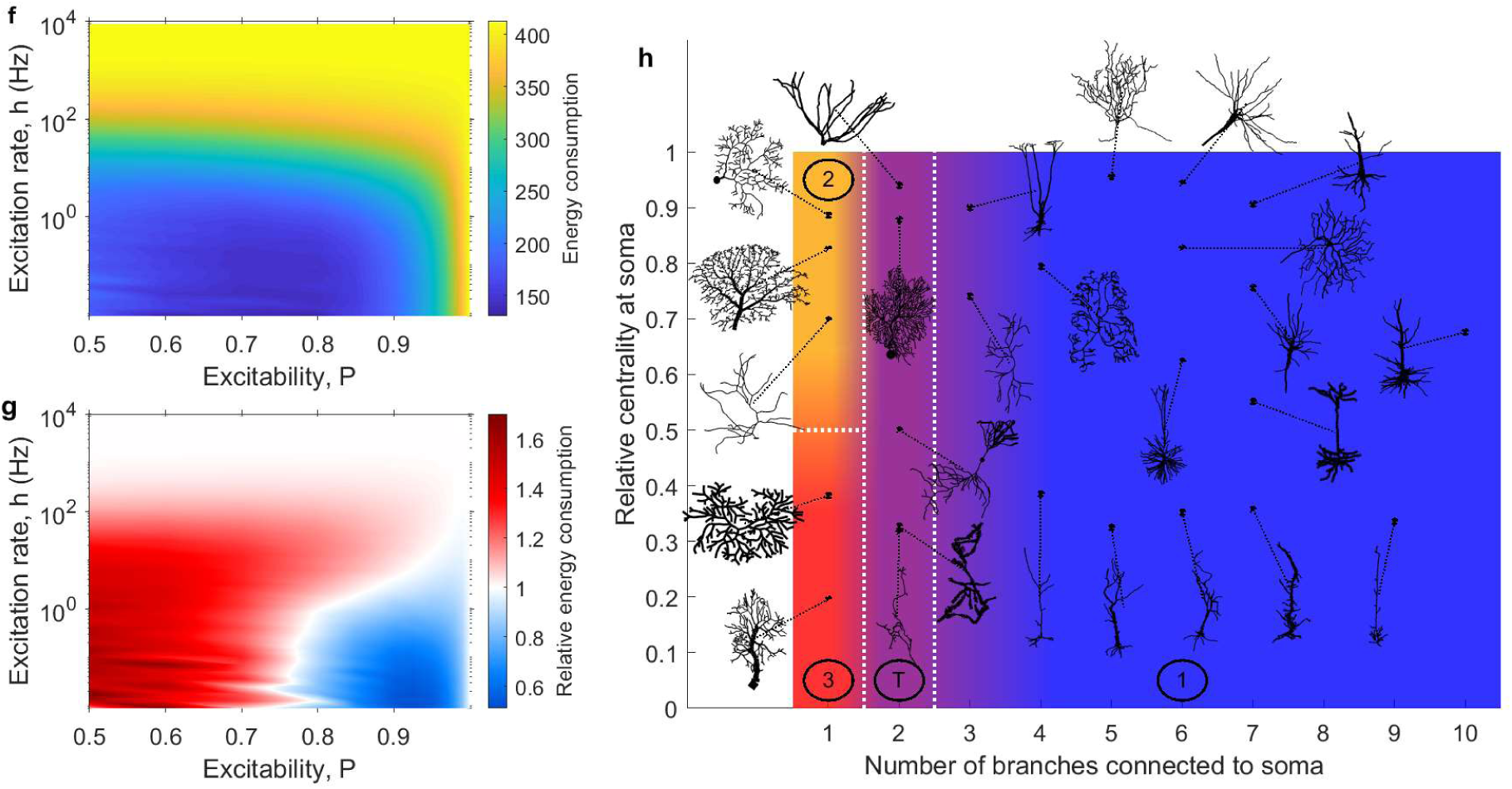
Dynamical description of the neuronal model. (a) We use detailed digital neuron reconstructions from NeuroMorpho.org. The dendrites are partitioned into fine compartments (pictured here is Neuron E, see Methods and Table 1). (b) The network of compartments forms an excitable tree to account for dendritic spikes. Each compartment undergoes the same type of dynamics. A temporal evolution of states of a compartment is shown here, corresponding to the susceptible (green), spiking (red), and refractory (yellow) stages of a dendritic spike. (c) Snapshot of neuronal dynamics with *h* = 0.5 and *P* = 0.96 using Neuron F (Methods). For a full illustration of the dynamics, please refer to Supplementary Video 1. (d) The firing rate of each compartment can be plotted for a particular set of parameters *h* and *P* (here, *h* = 1.12 and *P* = 0.9 using Neuron B), which govern the external driving and the dendritic excitability (see Methods for details), to obtain a map of most active regions within the neuron. Warm colors correspond to high firing rates, and cold colors to low firing rates. (e) Plotting the average firing rate of a particular compartment against the input rate *h* yields the response curve. The range of *h* values over which the firing rate varies most (between 10% and 90% of the maximum firing rate, *F*10 and *F*90) is called the dynamic range Δ. This example uses Neuron B with *P* = 0.9. (f) The energy consumption of a neuron is given by the number of total dendritic spikes per somatic spike, and is plotted over the parameter space. This example uses Neuron B. (g) The relative energy consumption represents the number of times a dendritic compartment fires, on average, per somatic spike. If it is less than 1, the neuron is energy efficient because dendrites effectively amplify the neuronal output (somatic spikes). If it is larger than 1, the neuron is considered energy inefficient. This example uses Neuron E. (h) Based on the number of branches connected to the soma and the centrality of the soma, neurons can be classified into four functional types: energy efficient (Type 1), partially efficient/inefficient (Type 2), energy inefficient (Type 3), and a transitional type (Type T) which can exhibit a mixture of the other types. See reference [29] for more details on the functional classification of neurons based on energy consumption.

The highly heterogeneous structure of the dendrites leads to complex non-linear interactions between signals. As the size and complexity of the dendrites reduces, the activity in the neuron becomes sparser. This occurs mainly because the rate of new dendritic spikes due to synaptic input increases with the number of compartments. This dependence of the neuronal size and activity also leads to an overall effect on the energy consumption of the neuron (Fig. 2 f). The energy consumption also depends on the main parameters of the model, as it is more pronounced for larger values of *h* and *P*. In general, smaller neurons are more economical, and hence our model predicts that the energy consumption of a neuron reduces as it ages (Fig. 3). This reduction in energy consumption with aging is particularly effective during the expensive dynamical regimes (large input rate *h* and excitability *P*).

**Figure 3:**
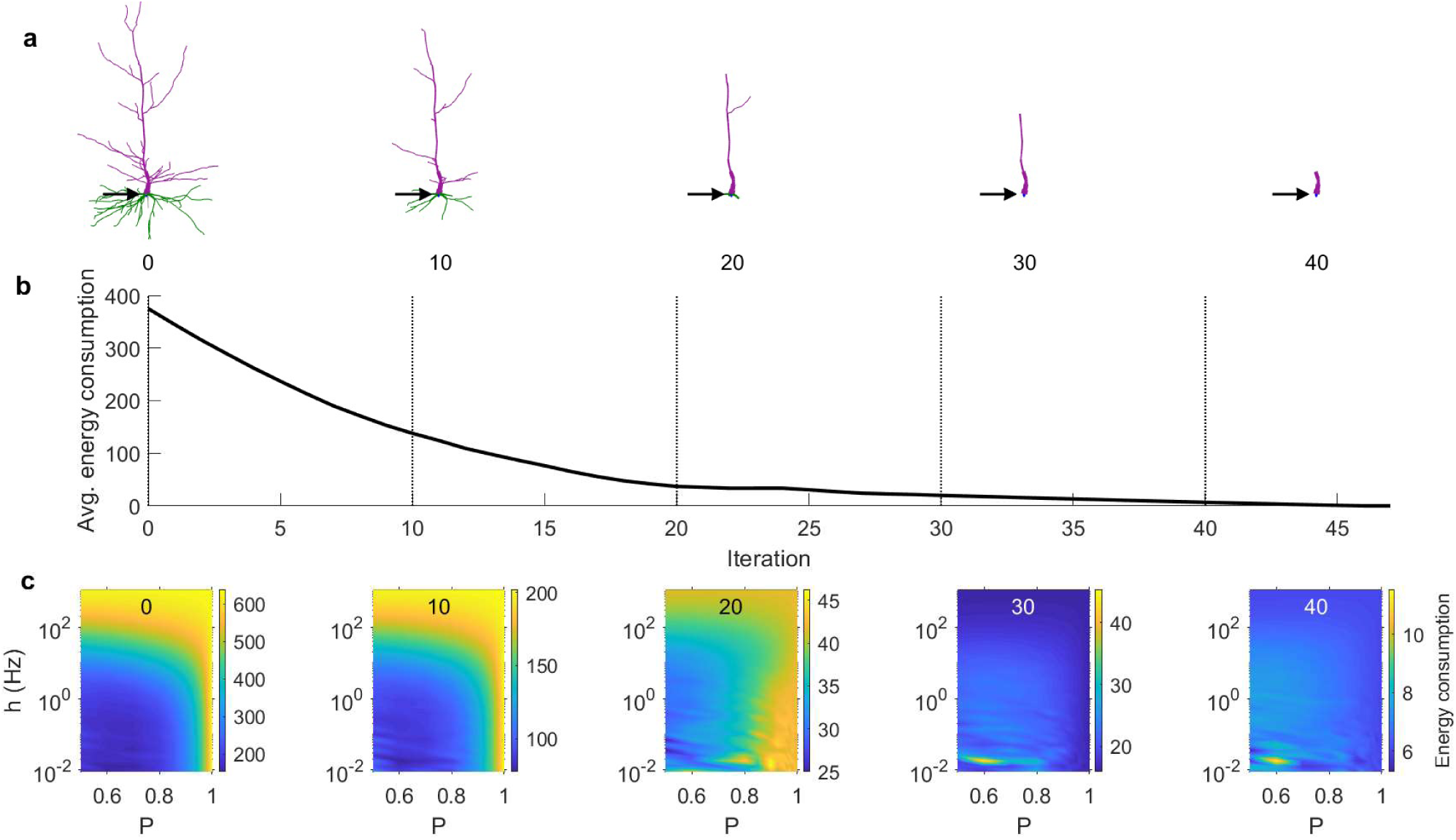
Effect of aging on total energy consumption. The total energy consumption is the total number of dendritic spikes per somatic spikes (see Methods for a formal definition). (a) Snapshots of the Neuron A after pruning iterations. (b) The total energy consumption of the neuron as a function of the pruning iteration, averaged over the parameter space 0.5 ≤ *P* ≤ 1 and 10^−2^ ≤ *h* ≤ 10^3^. (c) Snapshots of the total energy consumption at specific pruning iterations over a range of parameters *P* and *h*. The number at the top of each panel represents the iteration (marked by the vertical lines in panel **b**).

Another complementary measure is the relative energy consumption, which normalizes the energy consumption by the number of dendritic compartments, and compares the mean activity of dendritic compartments against the soma (see Methods). This relative energy consumption is inversely related to how efficiently the neuron uses its dendrites to fire. A low relative energy consumption is observed when the mean number of somatic spikes is larger than dendritic spikes. In this case, the neuronal topology amplifies the activity at the soma. As shown in Fig. 4, the relative energy consumption increases significantly as the complexity of the dendritic tree decreases. However, once all dendrites except the main apical shaft have been removed (see Fig. 1), it tends to decrease slightly. This highlights the importance of dendritic topology to shape the relative energy consumption. If the relative energy consumption is below 1, the neuron is generally efficient. If it is above 1, the neuron is generally inefficient. Therefore, as neuronal pruning accumulates, although the total number of dendritic spikes reduces, it becomes less energy efficient.

**Figure 4:**
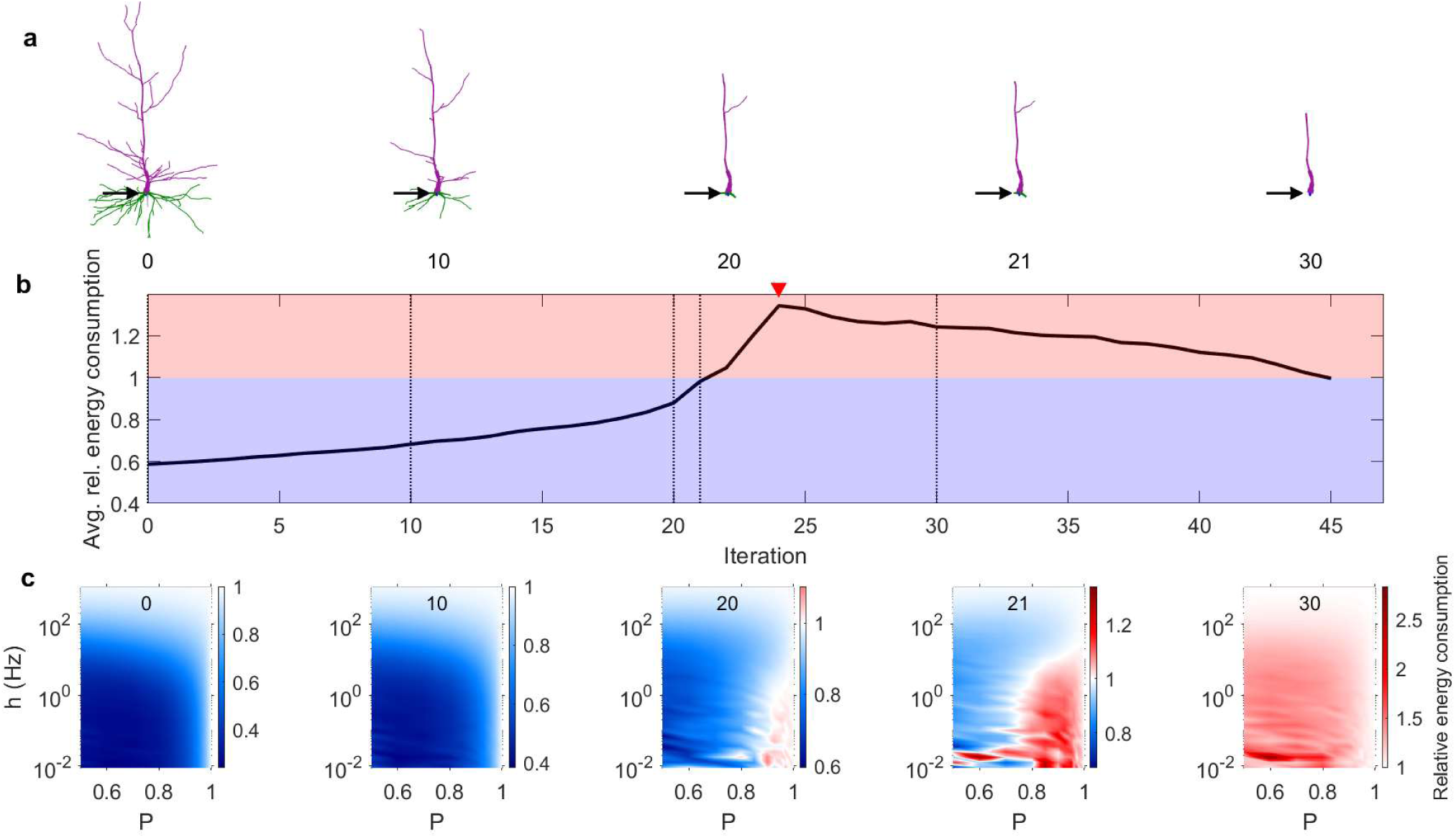
Effect of aging on relative energy consumption. The relative energy consumption is the mean number of dendritic spikes (across dendritic compartments) per number of somatic spikes (see Methods for a formal definition). (a) Snapshots of the Neuron A after pruning iterations. (b) The relative energy consumption of the neuron as a function of the pruning iteration, averaged over the parameter space 0.5 ≤ *P* ≤ 1 and 10^−2^ ≤ *h* ≤ 10^3^. At the red arrow, a single dendritic branch remains, and it corresponds to the most inefficient structure. (c) Snapshots of the relative energy consumption at specific pruning iterations over a range of parameters *P* and *h*. The number at the top of each panel represents the iteration (marked by the vertical lines in panel **b**).

Large dendritic arbors are known to increase the dynamic range [30], which determines what range of input strength the neuron can effectively encode in its output [31], and it is quantified based on the response function. Response functions reveal how the firing rate (output) of the soma changes with the input strength. As shown in Fig. 5, response functions are significantly affected by dendritic pruning. In general, pruning reduces the firing rate and the capability of the neuron to respond to weak stimuli. Therefore, the dynamic range is also progressively reduced with pruning iterations. The higher the signal propagation probability *P*, the higher the dynamic range. At high values of *P*, the dynamic range decreases more evenly over the pruning iterations. At lower values of *P*, the decrease in dynamic range occurs abruptly as the neuron approaches the point of major structural change (red arrow). It indicates the iteration at which only a single dendritic stem is connected to the soma.

**Figure 5:**
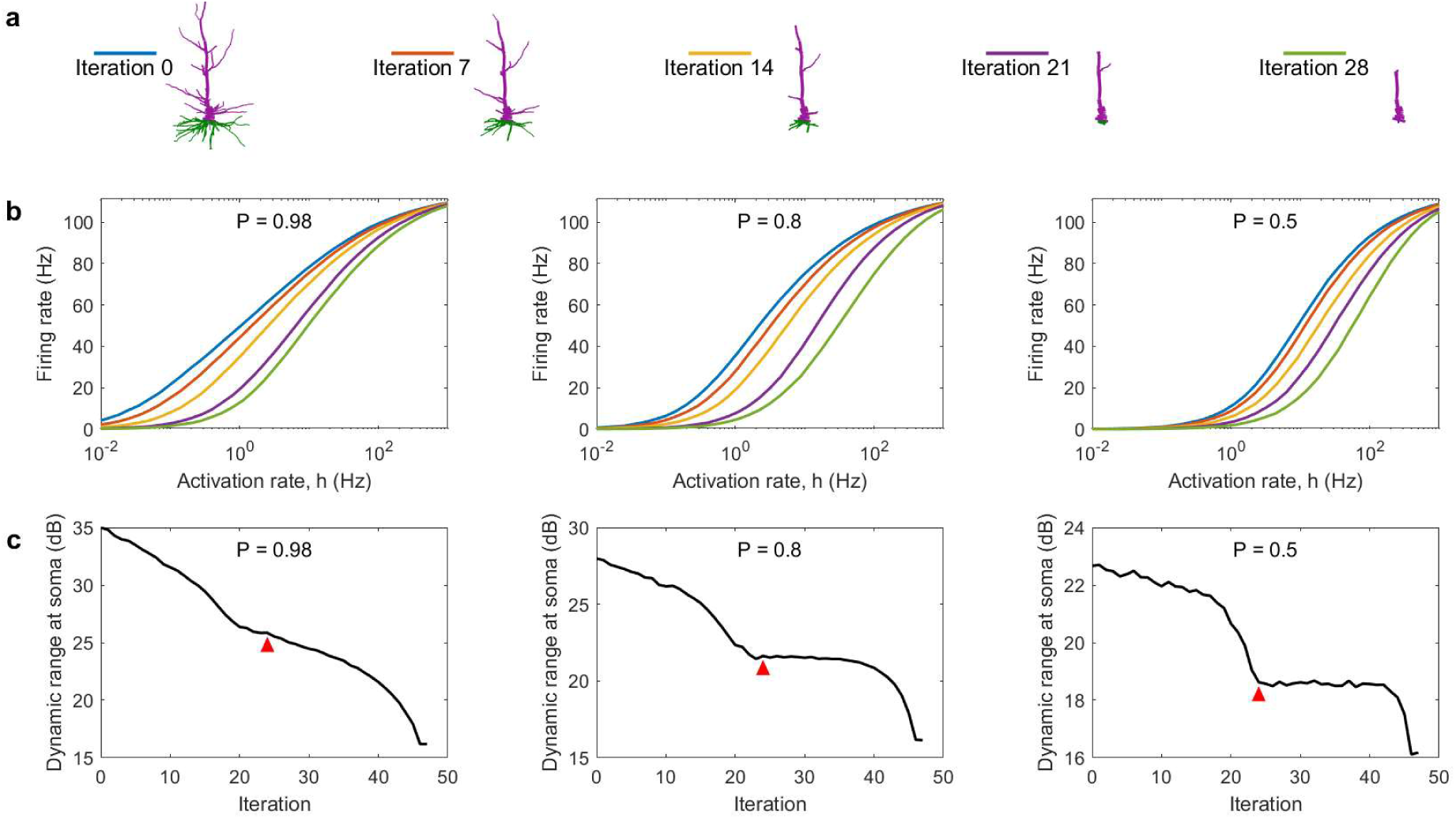
Change in dynamic range as a result of dendritic pruning. (a) Depiction of Neuron A at different pruning iterations. (b) Response function of the soma at various iterations corresponding to the plots in **a**. (c) Dynamic range at the soma for various *P* values. The red arrow indicates the point at which the number of somatic branches reaches 1 (iteration 24). Please note the different scales.

We have shown that Neuron A reduces its energy consumption as it ages, but becomes less efficient. Furthermore, the dynamic range at the soma is reduced with age. Fig. 6 shows that these findings also hold for other human pyramidal neurons. In each case, the point at which the number of somatic stems reduces to 1 (red arrow) closely corresponds to the most inefficient state (Fig. 6c).

**Figure 6:**
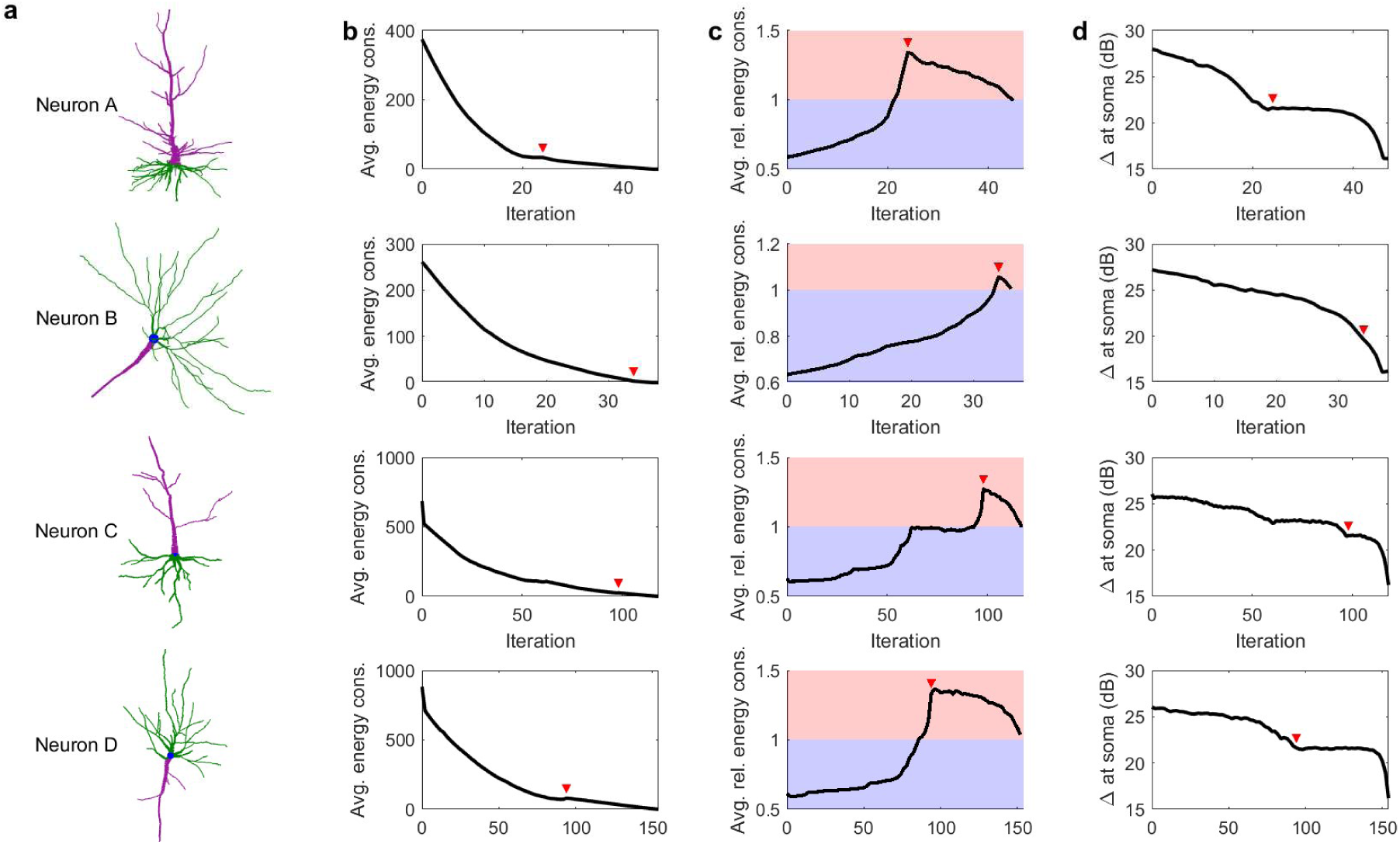
Effect of aging in the dynamics of different human pyramidal neurons. See Methods and Table 1 for a detailed description of the neurons. (a) Digital recreation of intact neurons. The soma is shown in blue, basal dendrites in green, and apical dendrites in purple. (b) The total energy consumption of the neurons as a function of the pruning iteration, averaged over the parameter space 0.5 ≤ *P* ≤ 1 and 10^−2^ ≤ *h* ≤ 10^4^. (c) The relative energy consumption of the neuron as a function of the pruning iteration, averaged over the parameter space 0.5 ≤ *P* ≤ 1 and 10^−2^ ≤ *h* ≤ 10^3^. Energy efficient stages are represented by the blue region, while energy inefficient stages are represented by the red region. (d) Dynamic range at the soma for *P* = 0.8. The red arrows mark the critical point at which the number of somatic branches reaches 1.

In a previous work [29], we proposed a functional classification of different types of neurons based on their relative energy consumption. Neurons were grouped into four categories depending on the relative centrality of the soma, and the number of dendritic branches connected to the soma. Type 1 indicates energy efficient neurons and have multiple branches connected to the soma; Type 2 neurons and Type T have both efficient and inefficient regions but Type 2 has only one branch connected to the soma while Type T has bitufted neurons; and Type 3 neurons are inefficient. This classification can be used to explore the relationship between changes in neuronal topology that takes place with pruning and energy efficiency. We subjected neurons to aging, and mapped them into the classification space according to the structural changes that occur (Fig. 7). As aging progresses, the number of dendritic branches and the relative centrality of the soma decrease monotonically. Increasing pruning, both centrality and the number of branches connected to the soma reduce. In this process that takes place with age, energy efficient neurons of Type 1 approach and cross to the transition Type T and then to Type 3, which is inefficient. Crucially, the process is not homogeneous as some pruning steps yield large jumps in this parameter space whereas others are essentially harmless.

**Figure 7:**
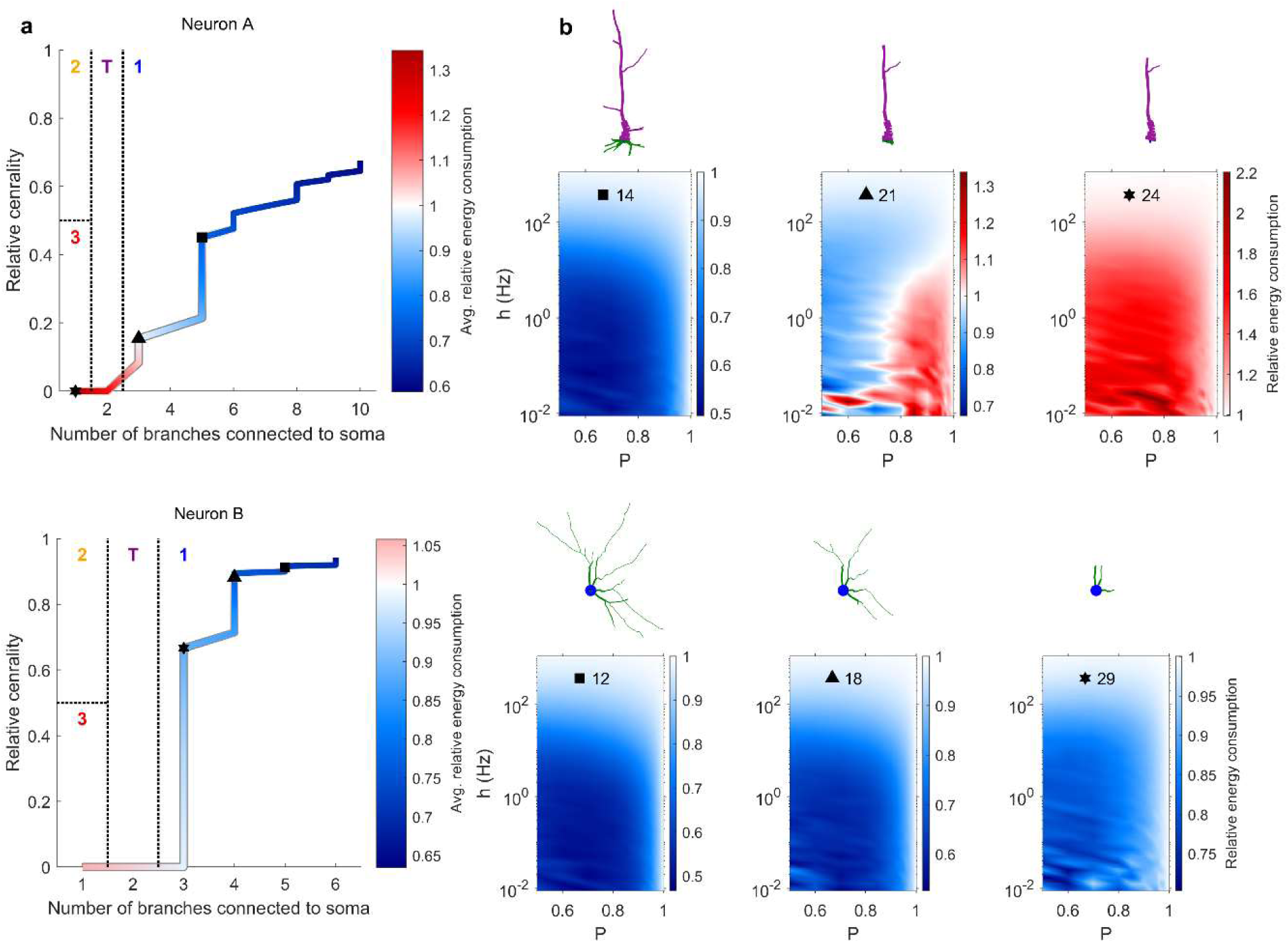
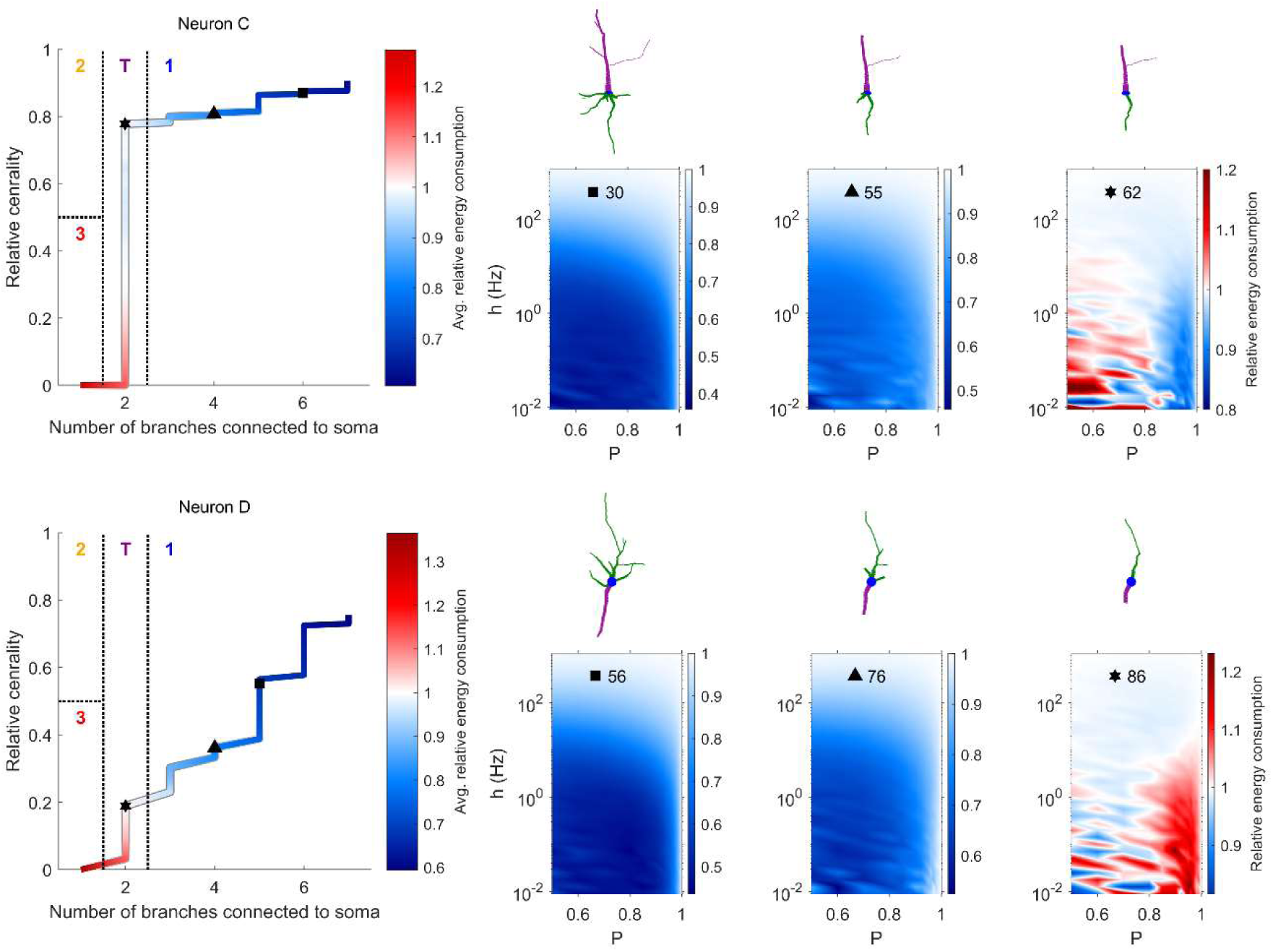
Neuronal classification scheme varies with pruning. (a) Evolution of four human neurons in the branch-centrality space that is used for classification. Color corresponds to the relative energy consumption, averaged over the parameter space 0.5 ≤ *P* ≤ 1 and 10^−2^ ≤ *h* ≤ 10^3^. The three symbols (square, triangle, and star) indicate the points at which snapshots are shown in panel **b**. (b) Snapshots of the dendritic structure and corresponding relative energy consumption profile after the specified number of pruning iterations.

## Discussion

Aging in the brain can have adverse effects on memory and normal cognitive function. Similar impairments can also be observed in neurodegenerative diseases such as dementia and Alzheimer’s disease [11, 21]. Understanding how dendritic recession contributes to this can be an important step in the mitigation and prevention of some of the negative effects. Here we put forward a model of dendritic pruning to study the dynamical and functional effects of aging and neurodegeneration at the neuronal level. We find that dendritic pruning reduces the total energy consumption. However, despite that, the neuron also becomes less efficient, as it usually requires more dendritic spikes to generate a neuronal spike. Moreover, this pruning reduces the dynamic range – the neuron’s ability to distinguish incoming input that varies over several orders of magnitude.

There is a notion of “successful” aging, in which the impairments caused by age are minimal [6, 32]. What are the criteria under which a neuron may continue to function normally while undergoing dendritic recession? Our results suggest that the centrality of the soma is a main indicator of the neuron’s resilience to aging, or a “*neuronal reserve”*, as an analogy to brain reserve [27], neural reserve [33-35], and cognitive reserve [28]. For example, neurons B and C have somas that are located centrally (Fig. 7). The more central a soma is located, the more evenly the dendrites are pruned towards the soma, allowing it to maintain its high centrality over many pruning iterations. Neurons with central somas are able to achieve better efficiency of the dendritic tree (lower relative energy consumption). In contrast, a non-central soma results in an asymmetric pruning, which quickly puts the soma at an even less central position. This accelerated reduction in centrality is observed for neurons A and D (Fig. 7, see also Fig. 6c). Therefore, our results suggest that the centrality of the soma is crucial to preserve neuronal performance, and a main indicator of neuronal reserve (or maintenance [32]), which quantifies the neuronal resilience to aging.

Modeling studies with neurons with idealized and regular tree structure showed that the size of dendritic trees plays a major role to increase the dynamic range [36]. A prediction from these regular Cayley trees was that large dendrites amplify weak incoming stimuli and the dynamic range [30]. Our results are consistent with these predictions. Furthermore, previous work concluded that bifurcations are an important topological feature that can increase the dynamic range [29]. Consistently with these results, a reduction in size and complexity of dendrites implies a lower dynamic range. However, our results also show that the dendritic pruning affects the dynamic range non-homogenously. Pyramidal neurons can eventually reduce to a single-branch neuron during their aging process. This is a landmark topology that affects the dynamic range and energy consumption of neurons. A single-branch neuron is a one-dimensional network of coupled active units, for which analytical approximations are known [37]. For example, the approximations predict that the dynamic range is enhanced for higher values of *P*, which is in agreement with our results (Fig. 4). For the multi-dimensional case, mean-field approximations also exist to determine the maximum dynamic range a system can attain [38]. Our results demonstrate that the dynamic range of young neurons decreases as more dendritic compartments are removed. In contrast, the dynamic range of aged (nearly one-dimensional) neurons remains essentially stable for lower values of *P* (Fig. 5).

Currently, the timescales involved in the dendritic recession in aging neurons are not known. Here, we have made the assumption that the dendrites are pruned progressively and evenly. Over time, this successfully reproduces morphologies with reduced complexity and size that have been observed experimentally (Fig. 1). In the future, our algorithm can be modified to stochastically remove *N* terminal compartments per iteration. Furthermore, it has also been suggested that aging neurons can lose whole branches at a time [13]. This could be implemented by removing internal compartments and their children. By observing the severity in the change of dynamic range caused by such a removal, it may be possible to identify and classify dendritic segments that are critical in maintaining the proper functioning of the neuron.

Functional decline in working memory that takes place during aging can be associated to the function of specific neuron types [1]. Cognitive studies of human subjects have revealed that memory degradation can especially inhibit the ability to discern similar experiences, called pattern separation [39, 40]. Our results suggest a neural basis of these behavioral effects of aging. If we assume that similar patterns correspond to similar rates of synaptic input, we can compare the neuronal output for these values of input of an aged neuron to a neuron whose dendritic integrity is intact (Fig. 8a). Because young neurons have larger dynamic range, for slight variations in relatively weak stimuli, we can see that dendritic pruning adversely affects the neuron’s ability to discern changes of input strength, which can lead to impairment in pattern separation, or the ability to distinguish between the different input values.

**Figure 8:**
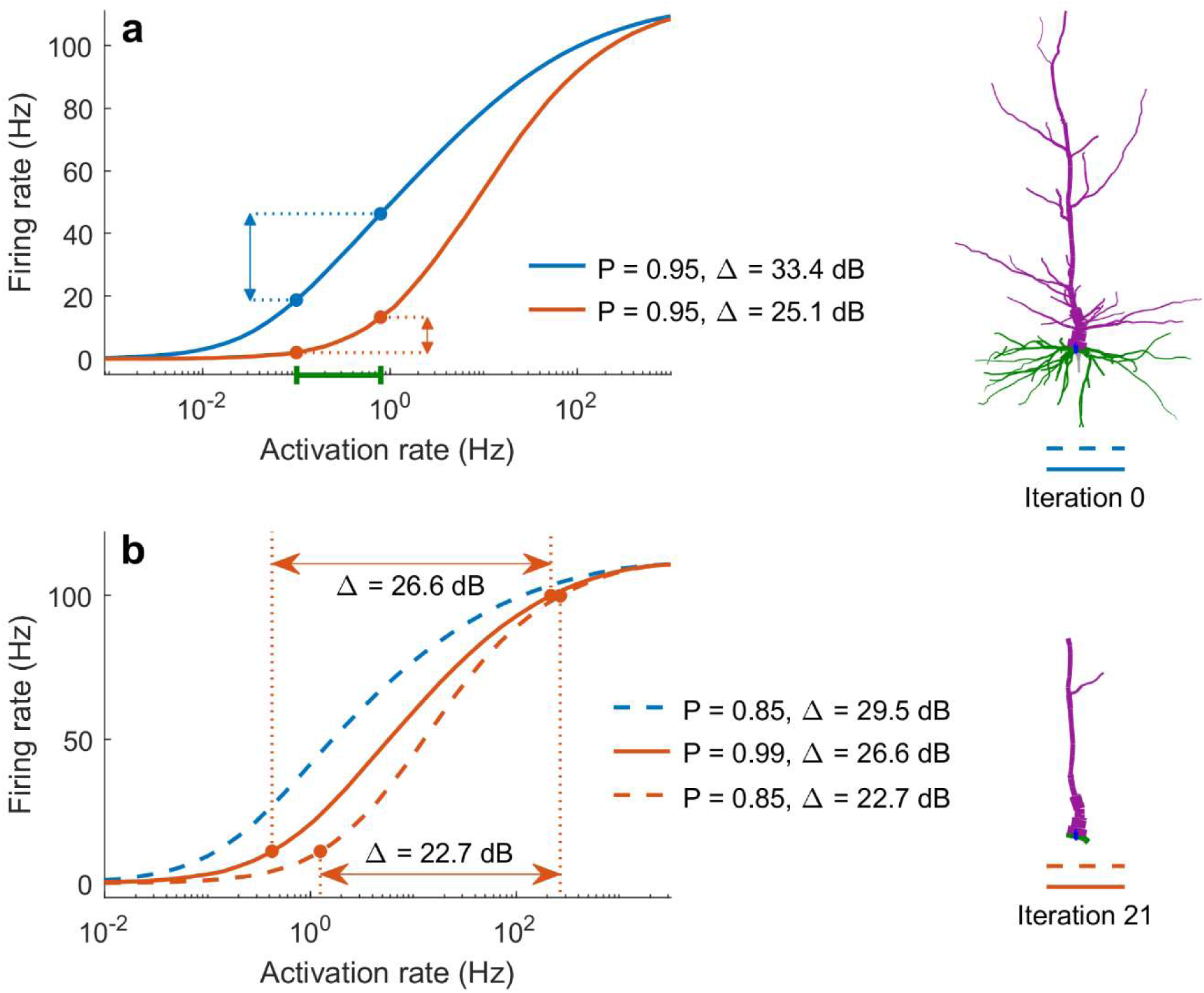
Effects of aging on the neuronal dynamic range. (a) The response functions of a young (blue) and aged (orange) neuron reveal that young neurons can more effectively distinguish and encode similar strengths of input (green) because they have a larger dynamic range. Here, *P = 0.95*. (b) Aging-related pruning decreases the dynamic range of neurons. By increasing the dendritic excitability, *P*, ageing neurons can partially mitigate this effect (solid orange line).

Moreover, it has been shown in primates that by manipulating potassium channels, it is possible to partially restore the physiological integrity of neurons [1]. Following a similar approach, our results indicate that excitability could also be explored to mitigate reduced cognitive abilities in elderly. In particular, the dynamical effects of aging on the response function and dynamic range can be at least partially restored by increasing the neuronal excitability *P* (Fig. 8b). Since this parameter in our model is rather general and unspecific, multiple possible electrophysiological manipulations could give rise to this net effect over *P*. Although further work is required, these ideas of rejuvenating neuronal dynamics can be a promising avenue of future research in an aging society.

## Methods

### Neurons and database

Detailed digital reconstructions of neurons were taken from the NeuroMorpho database [41-46]. The dendritic branches are made up of often thousands of connected compartments and include many bifurcations. Here, the soma is represented as a single compartment. The list of neuron reconstructions used here is given in Table 1.

**Table 1:** List of neuron reconstructions used in this study, taken from NeuroMorpho.Org (version 7.6).

| Label | Cell type | Species | Number of somatic branches | Number of compartments | Number of bifurcations | NeuroMorpho ID | Reference |
| --- | --- | --- | --- | --- | --- | --- | --- |
| A | Pyramidal | Human | 10 | 640 | 49 | NMO_84457 | [47] |
| B | Pyramidal | Human | 6 | 413 | 24 | NMO_03500 | [48] |
| C | Pyramidal | Human | 7 | 1091 | 222 | NMO_01064 | [49] |
| D | Pyramidal | Human | 7 | 1441 | 213 | NMO_01058 | [49] |
| E | Interneuron | Drosophila Melanogaster | 1 | 4659 | 762 | NMO_51008 | [50] |
| F | Purkinje | Mouse | 1 | 1568 | 129 | NMO_54509 | [51] |

### Pruning algorithm

It has been shown that the dendritic arbor in aging neurons can retract, decreasing both in size and complexity (branching order) [13, 15, 19]. For this study, we considered an iterative approach to simulate these changes. We start with the full neuron reconstruction, as provided by NeuroMorpho. A compartment that does not have any children is considered a terminus. In each iteration, we remove all terminal dendritic compartments. This has the effect of progressively pruning each branch towards the soma. Our main focus is on the effects of topology on neuronal dynamics. Since the neurodynamical model used here does not explicitly model synapses, synaptic changes relating to aging [6] are ignored. Similarly, changes in the shape or radius of dendritic branches [13], as well as chemical changes [52], are not explicitly modeled.

### Neuronal dynamics

Compartmental spikes were modeled using an excitable network of dendritic compartments and the somatic compartment, as seen in previous studies [30, 31, 53]. Compartments are modelled as probabilistic cellular automata with synchronous update [54]. A compartment in the susceptible state can be activated either by receiving the signal from a neighboring compartment with probability *P*, or by external input with rate *h* [29, 30, 55]. External inputs represent dendritic spikes invoked by synapses. Since not much data is available regarding the locations of synapses along the dendrites (there can be thousands), and since many underlying factors are responsible in the synaptic transmission process, the external input was modeled stochastically, with the probability of activation at some time step given by *r* = 1 − exp(−*h* × *δt*), where *δt* is the time resolution (0.001s). Active compartments will transition to the refractory state in the next time step, in which they remain for 8 steps, preventing sustained activity [56], then revert to the quiescent state. The three states correspond to the spiking behavior of electrically excited compartments (Figs. 2b, c).

Average compartmental firing rates as a function of *h* are used to construct the response function (illustrated in Fig. 2e). It maps the input-output behavior of the compartment [29]. A characteristic quantity derived from the response function is the dynamic range, defined as

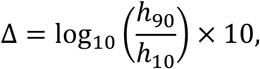

where *h*_10_ is the input rate required to produce *F*_10_, and *h*_90_ is the input rate required to produce *F*_90_. *F*_*x*_ is the firing rate that corresponds to *x*% of the maximum possible firing rate [31].

The simulations were run for a duration of 500,000 time steps on each pruning iteration of a given neuron. The activation rate *h* was varied over several orders of magnitudes, and the propagation probability *P* was varied over 0.5 ≤ *P* ≤ 1. We have neglected *P* < 0.5 because the signal dies out quickly in this regime.

### Energy consumption

Energy is required to control the movement of ions during a dendritic spike. Despite differences in behavior due to variations in chemical and geometric properties of the dendritic membrane, we generalize spikes such that a firing compartment consumes one unit of energy [29]. Therefore, the energy required to generate a spike at the soma is given by the ratio of the number of spikes in dendritic compartments to the number of somatic spikes during the course of the simulation:

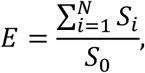

where *S*_*i*_ refers to the total number of spikes in compartment *i*, compartment 0 refers to the soma, and there are *N* dendritic compartments.

To gauge how effectively the neuron uses its dendrites to generate somatic spikes, we use a similar measure, called the relative energy consumption,

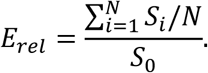

If *E*_*rel*_ < 1, dendritic compartments generally fire less often than the soma. Dendritic spikes are required only for integration [23], whereas somatic spikes are vital because they directly govern the functioning of the neuron [57, 58]. Thus, we say that the neuron is energy efficient. Conversely, if *E*_*rel*_ > 1, the neuron is energy inefficient.

### Centrality

A compartment’s centrality is defined as the distance (number of compartments) to the farthest terminal compartment, and is scaled between 0 and 1 relative to the centralities of all other compartments [29]. The most central compartment in a neuron has a centrality of 1, while the least central compartment has a centrality of 0.

### Code availability

MATLAB code for simulating neurodynamics and dendritic pruning for any neuron in the NeuroMorpho database is available at http://www.sng.org.au/Downloads.

